# Developing a primary *Paralichthys olivaceus* gill epithelial cells as an *in vitro* model for propagation of VHSV show a corresponding increase in cell viability with increase in protein concentration in growth media

**DOI:** 10.1101/708446

**Authors:** Sori Han, Jimin Hong, Jong-pyo Seo, Hetron Mweemba Munang’andu, In-kyu Yeo, Sung-hyun Kim

## Abstract

**Background:** Viral Hemorrhagic Septicemia Virus (VHSV) is a rhabdovirus that causes high mortalities linked to high economic losses in aquaculture. It has been grouped in four genotypes of which some do not easily propagate on continuous cell lines. As an alternative, the objectives of this study was to develop a primary gill epithelial cell (GEC) model from olive flounder (*Paralichthys olivaceus*) as an in vitro model for the propagation of VHSV.

**Results:** Our findings show that the primary GECs developed herein are highly permissive to replication of the JF-09 genotype IVa strain leading to high cytopathic effect observed within 96 hours post virus inoculation. Our findings also show that the viability GECs produced herein corresponded with increase in the concentration fetal bovine serum in growth medium. We envision that GECs produced herein will heighten our understanding of immune mechanisms associated with virus entry on gill mucosal surfaces in flounder.

## Introduction

Primary epithelial cells are increasing being used to study host response to infection because they provide insights on host pathogen interactions at portal of virus entry on mucosal surfaces. They provide vital information on cellular properties associated with susceptibility such as surface receptors that bind viral epitopes linked to infection establishment as we as a key data on immunological responses at portals of virus entry. Contrary to permanent cell lines, primary cells maintain vital cellular properties that favor virus replication essential for bulk antigen production in vaccine development (Alge et al., 2006; Pan et al., 2009). Hence development of primary epithelial cells is important for diagnostic purposes as well as production of bulk antigens required for vaccine production.

Viral hemorrhagic septicemia virus (VHSV) is a -ssRNA *Novirhabdovirus* belonging to the family *Rhabdoviridae* (Ammayappa and Vakharia, 2009). In East Asia, VHS disease was first reported with mass mortality in olive flounder (*Paralichthys olivaceus*) in Japan in 1996. The virus was later found to be widely distributed in wild and farmed olive flounder in South Korea (Kim et al., 2009) causing high mortality linked to severe economic losses. It is divided in four genotypes (I-IV) with different sublineages (Tafalla et al., 2008; Tafalla et al., 1998) render primary gill epithelial cells (GECs) to be a better alternative. However, the majority of VHSV replication GECs have mostly been carried out using European and North America strains on rainbow trout (RBT) GECs. Brudeseth et al (Brudeseth et al., 2008) propagated the genotype-Ia from Denmark in RBT GECs while Pham et al (Pham et al., 2013) produced infectious progeny on RBT GECs from genotype IVb, which are North American VHSV strains. To our knowledge there are no studies carried out on GECs generated from Asian fish species. Hence, our objective was to develop primary GECs from olive flounder essential for evaluating VHSV infectivity *in vitro* as a reliable model to study innate immune responses in a primary barrier against VHSV infections (Kim et al., 2014).

## Materials and Methods

### Fish examination

Juvenile olive flounder (<15 g) were obtained from a farm in Jeju, South Korea. The body surface and gills were examined for signs of disease infection and deformities. Fish were examined for presence of bacterial pathogens by isolation on basic growth media such as a blood agar and brain heart agar (BHI). Given that Edwardsiella infection is endemic in Olive flounder on Jeju Island, fsih were also screened for Edwardsiella infection based on the method described by Han et al. (Han et al., 2017). In addition, selective media such as shigella salmonella agar and TCBS agar were used to Edwardsiella screening. Direct RT-PCR (HelixAmp^tm^) and Direct PCR (helixAmp^tm^) kit was used to screen for viruses like VHSV, red sea bream Iridovirus (RSIV), viral nervous necrosis virus (VNNV), marine birnavirus (M BV) and Hirame rhabdovirus (HRV) as described by (Cho et al., 2008). All fish used in the study had no viral and bacterial infection detected.

### Primary cell culture of gill epithelial cells

Primary GECs were isolated and cultured using a protocol modified from Kim et al (Kim et al., 2014). First, gills were dissected and disinfected using antibiotics (Gentamicin, Gibco; 10 mg/ml) and antifungal drugs (Amphotericin B, Gibco; 250 µg/ml). Blood clots in the gill filaments were removed, and the gills were trypsinized in 0.5%Trypsin-EDTA (Gibco) in a vortex for 15 min. Trypsin activity was stopped by adding 10% fetal bovine serum (FBS; Gibco) in phosphate-buffered saline (PBS). GECs were collected and seeded in cell culture flasks in Gibco Leibovitz’s L-15 Medium (Gibco) containing 10% fetal bovine serum (FBS) and 1% of gentamicin (10 mg/ml). Thereafter, the flasks were incubated at 20°C for 24 hrs. After 24 hrs, the suspended particles and blood cells in the cell culture medium were removed by washing thrice using PBS. Cell culture medium was replaced every 3 - 4 day until virus infection. The cells were observed under a microscope after fixing with 10% (v/v) formaldehyde (in PBS) for 30 min at room temperature. Primary GECs were observed under the microscope at 24 hrs and 96 hrs post-seeding.

### MTT viability assay

Cell viability was measured in CellTiter AQ_ueous_ one solution (Promega, USA). To determine the optimal concentration of FBS required for GEC growth, FBS was constituted at 5%, 10% and 20% concentrations in L-15 medium having 1% gentamicin (10 mg/ml). Cells (1×10^4^/well) were seeded in 0.2 ml L-15 medium/well in 96-well plates. After removing the medium, each the three FBS concentrations was added to five wells. On days 3 and 7 after incubation at 20°C, 10 µl of CellTiter AQ_ueous_ One solution was added to each well. The plates were incubated for 24 hrs in 20°C. Absorbance was measured at 540 nm using the EPOCH spectrophotometer (BioTek).

### Viral hemorrhagic septicemia virus infection of primary gill epithelial cells

To determine the susceptibility of the olive flounder primary GECs to VHSV infection, first passage cells were seeded into 24-well plates at a concentration 1×10^5^ cells per well. When the cells were confluent after 96h post-seeding, they were infected by the wild type VHSV (JF-09 genotype IVa) previously isolated from olive flounder (Kim et al., 2014) at a multiplicity of infection (MOI) of 1. Virus titration to determine the MOI was performed in *Epithelioma papulosum cyprinid* (EPC) cells using the 50% tissue culture infective dose (TCID_50_/ml) method previously described by Spearman - Karber method. After infection, the GECs were incubated at 20°C for 96 hrs. Cytopathic effect (CPE) was confirmed by microscopy after fixing the cells with 10% (v/v) formaldehyde in PBS for 30 min at room temperature.

To determine virus propagation in the olive flounder primary GECs, cells were seeded (1×10^4^) in 96-well plates. When CPE appeared after VHSV inoculation, the supernatant was collected for VHSV verification by PCR. Reverse transcription (RT) was performed using Direct RT-PCR kit (HelixAmp^tm^) according to the manufacturer’s instruction. VGsense (5’-CCAGCTCAACTCAGGTGTCC-3’) and VGanti (5’-GTCACYGTGCATGCCATTGT3’) primers were used for amplification of a 587 base region of the VHSV G gene (Nishizawa et al., 2002). 5 µl of sample and D.W. up to total 50 µl were added to RT-PCR reaction mix. PCR conditions were as follows; Pre-denaturation 95°C for 5 min, 40 cycles (94°C for 20 s, 52°C for s, and 72°C for 1 min) and post extension 72°C for 5 min. PCR products were subjected to electrophoresis using 1.0% agarose gels containing RedSafe^TM^ (iNtRon). They were visualized under UV light to detect the difference of virus concentration between the inoculated VHSV stock (post-infection 0hrs) and the supernatant of the well at post-infection 96hrs with band thickness.

## Results and Discussion

### Primary Gill epithelial cells

Figure 1A shows partial confluence of primary olive flounder GECs attained after incubation at 20°C for 24 hrs. The cells appeared healthy and adherent to the base of the cell-culture flasks. Figure 1B shows results of the MTT viability analysis in which the quantity of live cells corresponded with increase in the concentration of FBS added to the growth media as shown from the samples collected at 72 hrs and 168 hrs post seeding. Overall, the quantity of live cells was highest in 20% FBS at 72 hrs and 168 hrs post seeding being more than twice the number of viable cells observed in cells seeded with 5% FBS at 72 hrs and 168hrs post-seeding.

**Figure 1.**
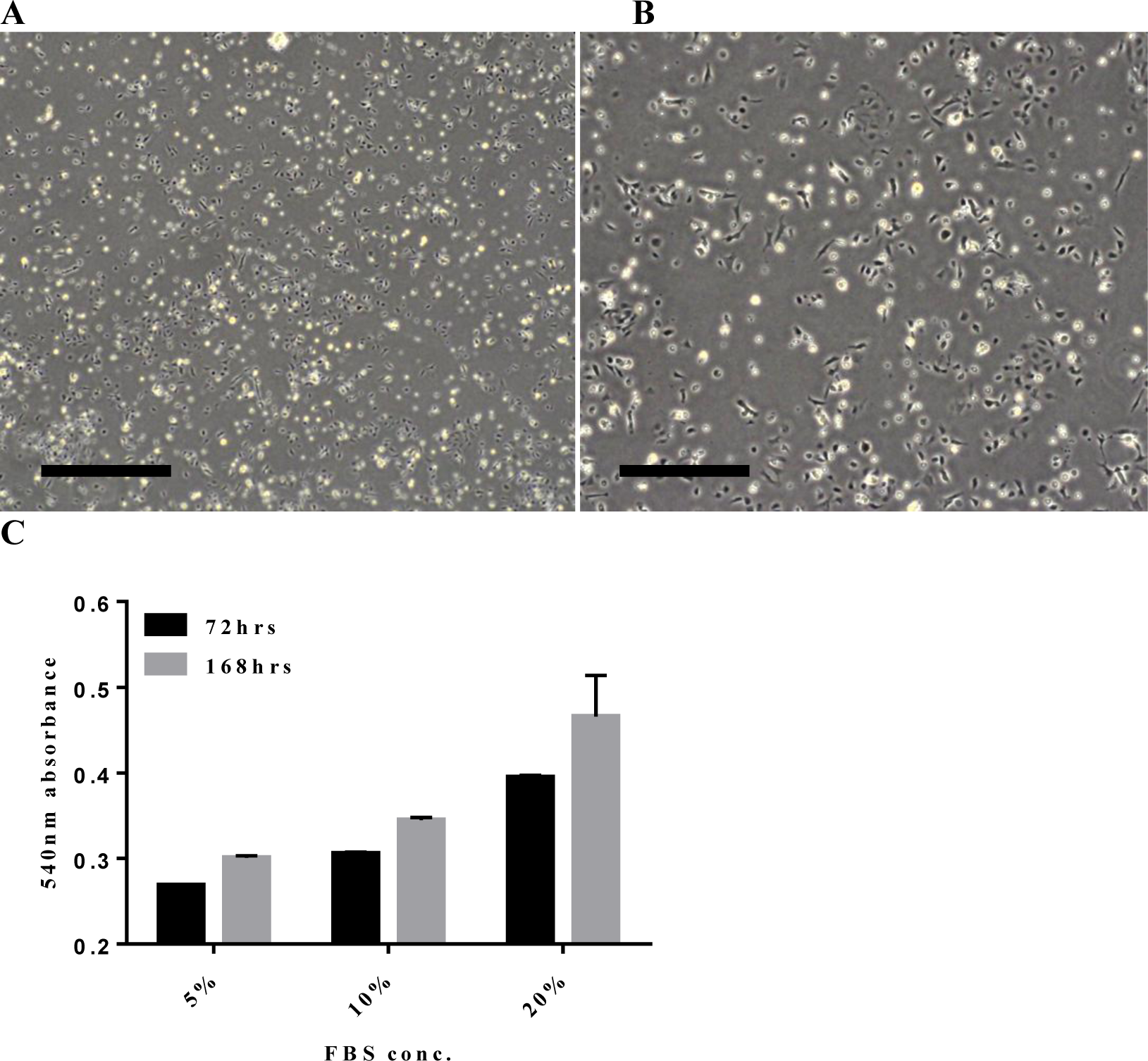
A shows a representative micrograph of primary GEC monolayer at 24 h post-seeding in L-15 medium supplemented with 20% fetal bovine serum (FBS). The micrographs were obtained using a light microscope (Olympus microscope CKX53) at 100x(1A) and 200x(1B) magnification. Bar = 500µm. Figure 1C: Primary GEC viability was measured using a CellTiter AQuous one solution reagent (promega) (N= 5) at post-seed 72hrs and 168hrs to show the viability and growth rate.

### Virus infection

Figure 2B shows that Olive flounder GECs generated in this study were permissive to VHSV replication as shown from CPE produced at 96 hrs and 120 hrs post virus inoculation. On the contrary, there was no CPE observed in the none-infected cells as shown in Figure 2A. Figure 3 shows detection of VHSV strain JF-09 genotype IVa infection by PCR. Note the detection of PCR products from GEC supernatants collected at 96 hrs post-infection shown by presence of positive bands in lanes 1, 2 and 3 that corresponded with detection of the VHSV in the positive control sample shown in lane P. On the other hand, there were no VHSV bands detected from uninfected GECs as shown in lanes 0_1_, 0_2_ and 0_3_ as well as the negative control distilled water (DW) in lane C. Put together, these findings demonstrate that the Olive flounder primary GECs developed in this study are permissive to VHSV strain JF-09 genotype IVa infection leading to production of CPE confirmed by PCR diagnosis.

**Figure 2.**
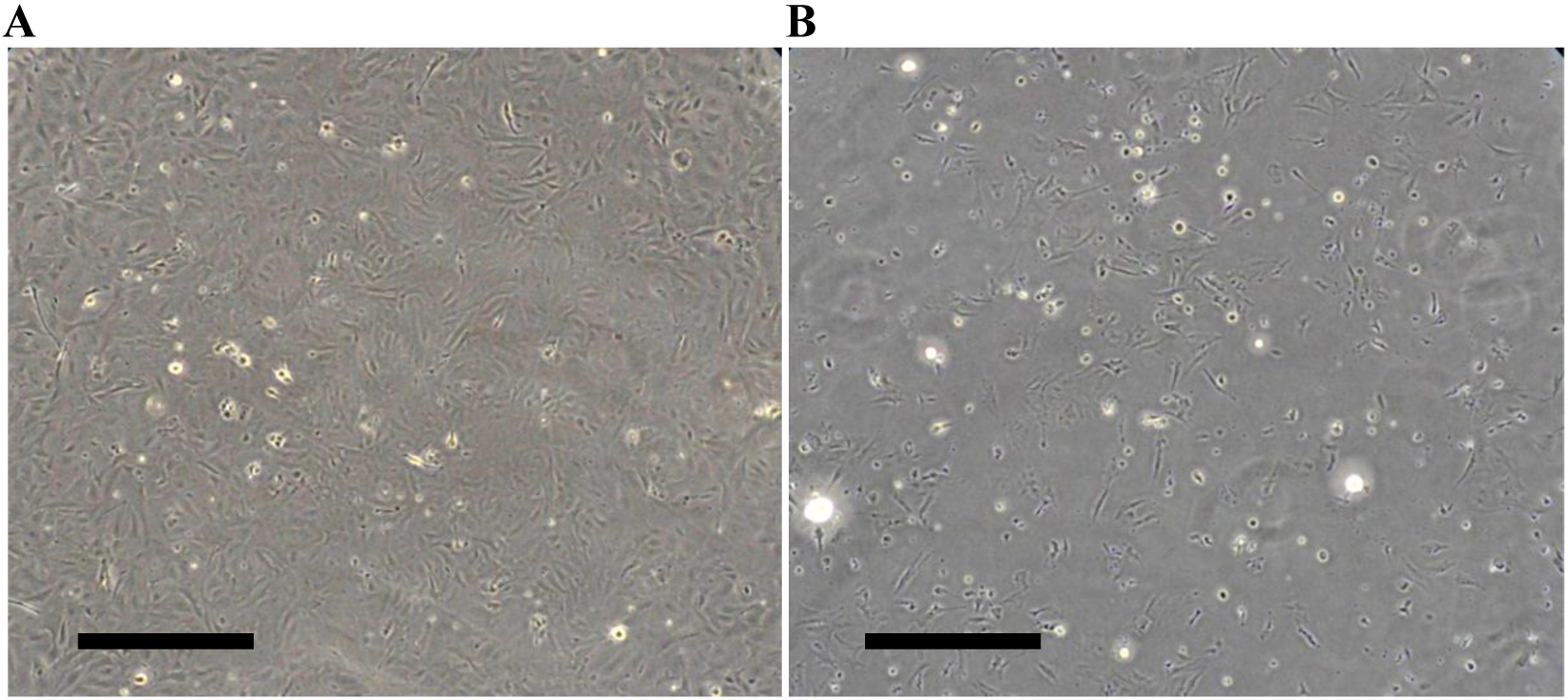
shows primary gill epithelial cells (GECs) monolayer generated from Olive flounder. Figure 2B shows development of cytopathic effect (CPE) 96 h post inoculation (hpi) of VHSV characterized by cell death. Figures 2A shows a none-infected GEC control. Micrographs were obtained using a light microscope (Olympus CKX53) at 100x magnification. Bar = 500µm.

**Figure 3:**
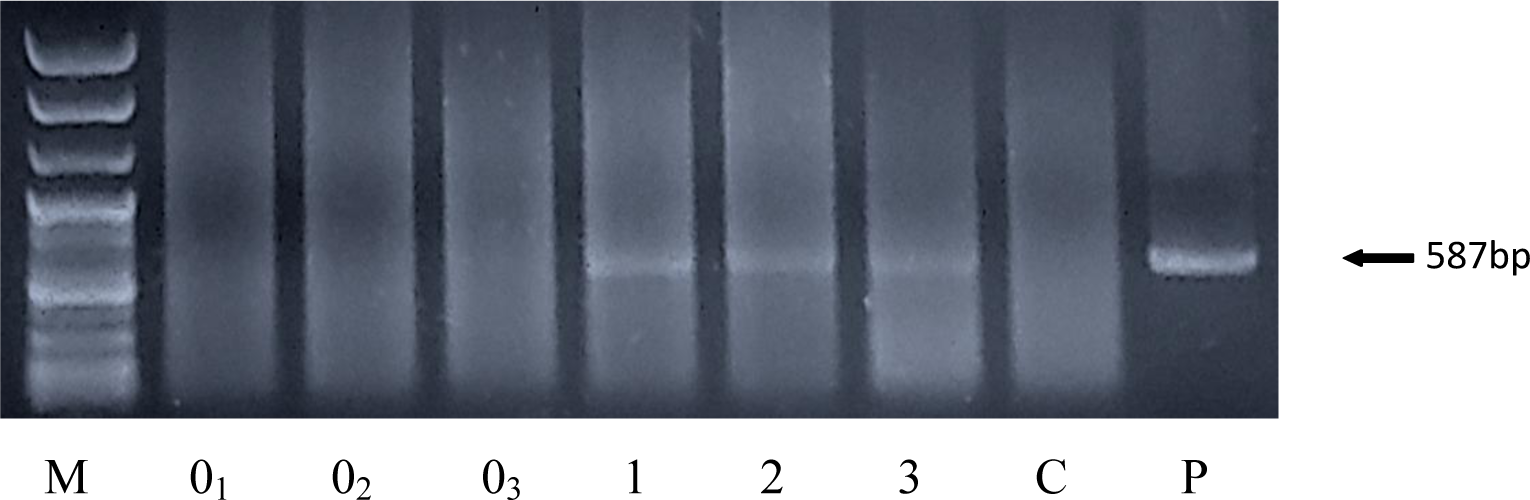
Virus replication was confirmed by RT-PCR using supernatants from GEC at 96hrs of incubation (MOI=1). 100bp DNA ladder maker (geneall): ‘M’; Post-infection 0hrs: ‘0_1_,0_2_,0_3_’; Post-infection 96hrs: ‘1,2,3’; Control (not viral infected): ‘C’; Positive control: ‘P’

Our findings are in line with studies carried out on RBT GECs (*Oncorhynchus mykiss*). Pham et al (Pham et al., 2013) showed that RBT gill epithelial cells were more susceptible to VHSV infection than spleen macrophages from because they led to rapid growth rate, higher CPE and virus yield. Tafalla et al (Tafalla et al., 2008) showed that VHSV was not able to complete its replication cycle in monocyte/macrophage-like cell line RTS11 and, hence, it failed to produce infectious viral particles. They attributed inhibition of the virus replication to high IFN and Mx expression by RTS11 cells. In another study, Tafalla et al (Tafalla et al., 1998) showed that VHSV infected trout and turbot head kidney macrophages as well as blood leukocytes primary cultures, but failed to produce CPE. Brudeseth et al (Brudeseth et al., 2008) showed that the virulent VHSV strain propagated on RBT GECs caused high CPE and translocated into neighboring cells within 2 h post inoculation (dpi) and yet on primary head kidney cells it only produced 9.5% maximum infectivity 3 dpi. Several studies have shown that cell-lines of lymphoid origin like macrophages and monocytes with high IFN expression inhibit the growth VHSV. Although we did not examine the IFN expression levels in this study, the general observation is that GECs are low IFN producers compared to macrophages and monocytes thus make good candidates for VHSV propagation. Hence, the approach used in this study would aid in diagnosis and vaccine development.

Hence, in the present study we inoculated the JF-09 genotype IVa on primary GECs generated from olive flounder resulting in a high replication rate and CPE formation within 72 h after inoculation clearly showing that the olive flounder primary GECs are highly permissive to rapid propagation of the JF-09 genotype IVa strain. Hence, in situation where rainbow trout are not available for preparation of GECs, olive flounder can be used to prepare GECs for the rapid propagation of VHSV with the view to obtain high virus yield using the approach used in this study. Moreover, our findings show that 20% FBS gives high cell viability than 5% and 10% indicating the method can be optimized to increase cell viability during propagation up to 168 hrs (1-week) post seeding. We envision that these findings shall contribute to development of diagnostic tools and vaccine production against VHSV strains infecting olive flounder and other fish species.

## Acknowledgements

We thank for all participants in this study

## Competing interests

The authors declare no competing or financial interests.

## Author contributions

Conceptualization: S.K., I.Y.; Methodology: S.K., S.H.; Formal analysis: S.H.; Investigation: S.H.; Resources: S.H., J.S., S.K.; Data curation: J.H.; Writing -original draft: J.H., S.H., S.K., I.Y.; Writing - review & editing: S.K., J.S., I.Y., H.M.M.; Supervision: S.K., H.M.M., I.Y.; Project administration: S.H., S.K.

## Funding

This work was supported by Korea Institute of Planning and Evaluation for Technology in Food, Agriculture and Forestry (IPET) through Golden Seed Project, funded by Ministry of Agriculture, Food and Rural Affairs (MAFRA). (213008-05-2-SB220).

